# TRGT-denovo: accurate detection of *de novo* tandem repeat mutations

**DOI:** 10.1101/2024.07.16.600745

**Authors:** T. Mokveld, E. Dolzhenko, H. Dashnow, T. J. Nicholas, T. Sasani, B. van der Sanden, B. Jadhav, B. Pedersen, Z. Kronenberg, A. Tucci, A. J. Sharp, A. R. Quinlan, C. Gilissen, A. Hoischen, M. A. Eberle

## Abstract

**Motivation:** Identifying *de novo* tandem repeat (TR) mutations on a genome-wide scale is essential for understanding genetic variability and its implications in rare diseases. While PacBio HiFi sequencing data enhances the accessibility of the genome’s TR regions for genotyping, simple *de novo* calling strategies often generate an excess of likely false positives, which can obscure true positive findings, particularly as the number of surveyed genomic regions increases.

**Results:** We developed TRGT-denovo, a computational method designed to accurately identify all types of *de novo* TR mutations—including expansions, contractions, and compositional changes— within family trios. TRGT-denovo directly interrogates read evidence, allowing for the detection of subtle variations often overlooked in variant call format (VCF) files. TRGT-denovo improves the precision and specificity of *de novo* mutation (DNM) identification, reducing the number of *de novo* candidates by an order of magnitude compared to genotype-based approaches. In our experiments involving eight rare disease trios previously studied TRGT-denovo correctly reclassified all false positive DNM candidates as true negatives. Using an expanded repeat catalog, it identified new candidates, of which 95% (19/20) were experimentally validated, demonstrating its effectiveness in minimizing likely false positives while maintaining high sensitivity for true discoveries.

**Availability and implementation:** Built in Rust, TRGT-denovo is available as source code and a pre-compiled Linux binary along with a user guide at: https://github.com/PacificBiosciences/trgt-denovo.

## Introduction

Tandem repeats (TRs) are DNA sequences composed of variably recurring, (nearly) identical subunits that contribute significantly to both intra-sample and population-level genomic variation [1]. *De novo* expansions of TRs, present in both coding and non-coding regions [2], are associated with over 60 monogenic disorders [3] and linked to conditions such as cancer [4,5] and neurological disorders [6]. The mutability of TRs, influenced by their repeat length and sequence context [7,8], is significantly higher—by several orders of magnitude—than that of non-repetitive DNA [8].

Accurately sizing large TR loci is challenging, specifically for pathogenic TR loci, whose size often expands significantly from normal to premutation to pathogenic repeat ranges [3]. The Tandem Repeat Genotyping Tool (TRGT) [9] was recently developed to characterize TR loci in PacBio HiFi sequencing data. TRGT calculates repeat length, composition, mosaicism, and CpG methylation state while also providing visualization. TRGT demonstrates a high Mendelian consistency rate exceeding 98.38% when excluding off-by-one errors, indicating its high accuracy. However, despite this high accuracy, the presence of millions of repeat loci in the genome means that trio analysis can still generate tens of thousands of false positives *de novo* calls. Thus, to integrate *de novo* TR analysis into rare disease studies effectively, a strategy is needed to filter out a substantial portion of these false positives while increasing specificity without missing true positives.

We present TRGT-denovo, a novel method for detecting DNMs in TR regions by integrating TRGT genotyping results with read-level data from family members. This approach significantly reduces the number of likely false positive *de novo* candidates compared to genotype-based *de novo* TR calling. In a follow-up to earlier research surveying DNMs in eight rare disease trios [10], we used the same data to demonstrate that TRGT-denovo would have accurately classified all high-quality candidate *de novo* calls—later experimentally validated as false positives—as true negatives. Moreover, by expanding *de novo* analysis using a larger repeat catalog in the same dataset, targeted sequencing confirmed 95% (19/20) of the selected *de novo* candidates detected by TRGT-denovo.

## Results

Throughout, we consider TR genotyping obtained by using TRGT in 9 family trios and the GRCh38 reference genome, along with various repeat catalogs [11,12]. TRGT-denovo performs *de novo* TR calling, as detailed in the methods section. Briefly, TRGT-denovo analyzes both the genotyping outcomes and reads spanning the TRs generated by TRGT, as shown in Fig. 1. TRGT-denovo compares the alleles and supporting reads from the child against those of the parents, enabling the identification and quantification of variations exclusive to the child’s data as potential DNMs. As a result, TRGT-denovo can detect both changes in TR length and compositional variations (e.g., sequence changes such as SNVs and larger).

**Fig 1.**
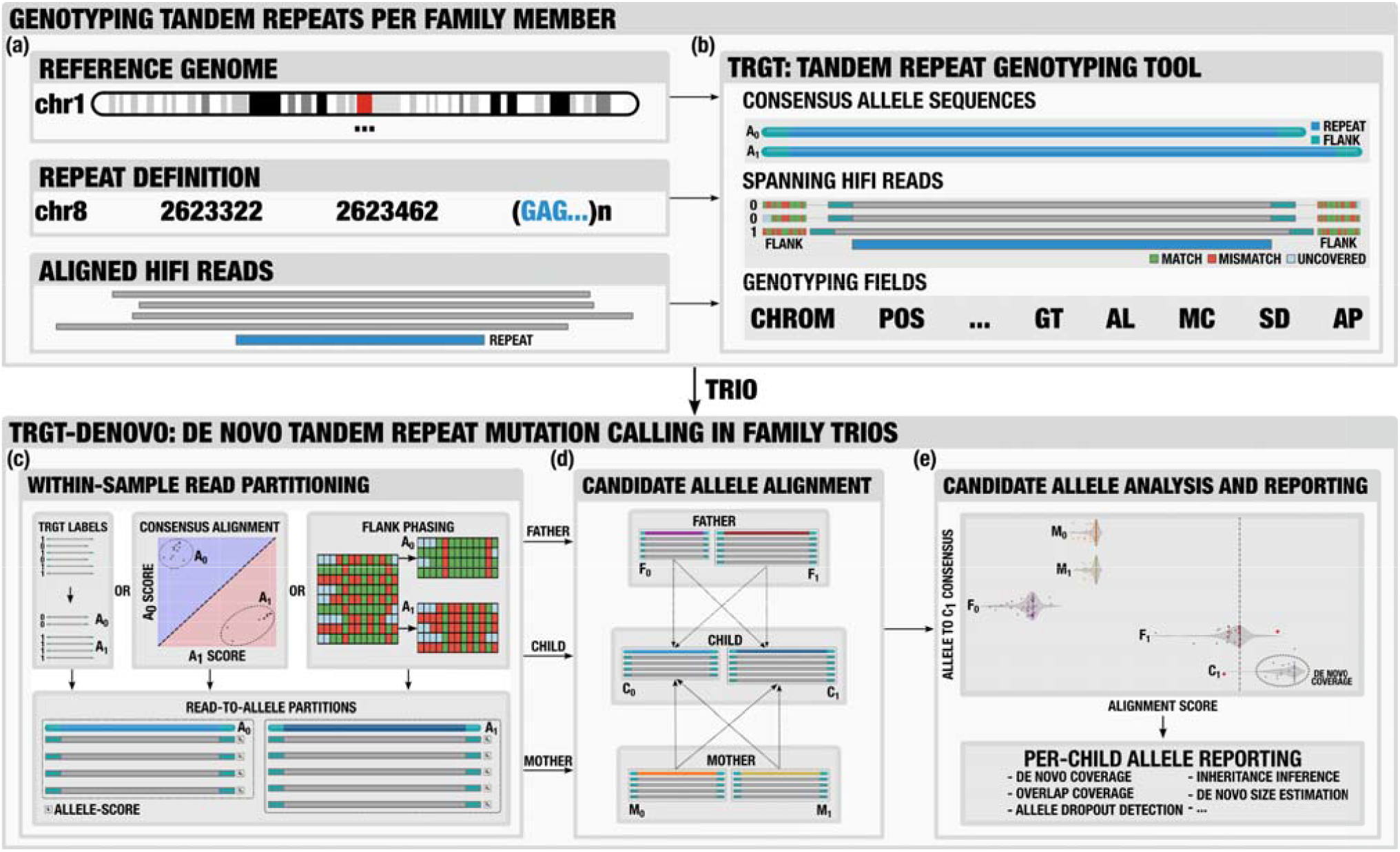
Overview of TRGT-denovo (full details in Methods). (a) TRGT pre-processing, which requires aligned PacBio HiFi reads, a repeat definition catalog, and a reference genome. (b) TRGT-denovo uses TRGT output, specifically spanning reads and genotyping data, along with the reference genome and repeat definitions. (c) By matching repeat definitions and corresponding allele sequences, reads are partitioned and assigned to alleles. This is achieved via TRGT-obtained classifications, consensus allele alignment, or phasing, thus determining the allele sequence each read best supports. (d) Allele partitioned reads are realigned to child allele consensus sequences for comparison purposes. (e) Potential DNMs are identified by examining discrepancies in alignment score distributions among candidate *de novo* alleles.

### Comparing genotype-based *de novo* TR calling to TRGT-denovo

We assessed *de novo* TR calling by conducting a comparative analysis on the HG002, HG003, and HG004 trio with 30x HiFi data sequenced on the PacBio Revio system. Using a repeat catalog of 891,328 loci (excluding those in segmental duplications), TRGT generated genotype calls across the trio at 888,711 (99.7%) loci. TRGT provides various metrics, including the repeat counts per allele and allele depth (number of reads spanning each allele) for each genotyped TR locus. Our analysis focused only on sites supported with a minimum of ten reads, with at least five reads supporting each allele in each trio member, resulting in 864,990 loci for evaluation. *De novo* candidates were identified by detecting deviations from expected Mendelian inheritance patterns, with 81,308 (9.40%) loci displaying Mendelian inheritance inconsistencies indicative of potential *de novo* mutations. Of these, 89.08% and 7.04% correspond to homopolymer and dinucleotide repeats, respectively. When allowing for variations of one motif count, 2,582 loci (0.29%) were flagged as potential *de novo* mutations, with 63.07% and 22.22% corresponding to homopolymer and dinucleotide repeats respectively.

TRGT-denovo enhances the detection of true *de novo* tandem repeat expansions, contractions, or compositional changes while reducing likely false positives by analyzing reads from all trio members (see Methods). In this dataset, TRGT-denovo found that 4,214 (0.49%) of analyzed loci showed some degree of *de novo* evidence when testing both alleles of the child. *De novo* evidence is defined as at least one read inconsistent with both parents’ called genotypes. The discrepancy between the 81,308 Mendel inheritance errors identified from genotype calls and the 4,214 loci showing *de novo* evidence stems from TRGT-denovo’s consideration of underlying read data in addition to inferred genotypes. Genotyping can be ambiguous; however, by analyzing read and allele length distributions within the context of a trio (i.e., comparing the child with both parents), we can get a more reliable indication of *de novo* mutations than by just counting motifs. Note that *de novo* evidence does not always indicate true DNMs; it could also result from artifacts like parental allele dropout, mosaicism, somatic instability, or stutter, particularly in low coverage scenarios. To address this uncertainty, TRGT-denovo assesses the amount of *de novo* evidence using several metrics, including the allele *de novo* ratio and the child *de novo* ratio, which respectively measure the proportion of *de novo* evidence within an individual allele and across the entire locus. Furthermore, the potential size of a *de novo* event is evaluated by the mean absolute difference between the *de novo* reads and the closest parental read data. This measurement provides insight into the event’s magnitude, with smaller values under low coverage conditions likely indicative of artifacts. Fig. 2. shows how *de novo* coverage (the number of reads supporting the *de novo* event) varies in relation to these metrics for alleles exhibiting *any de novo* coverage.

**Fig 2.**
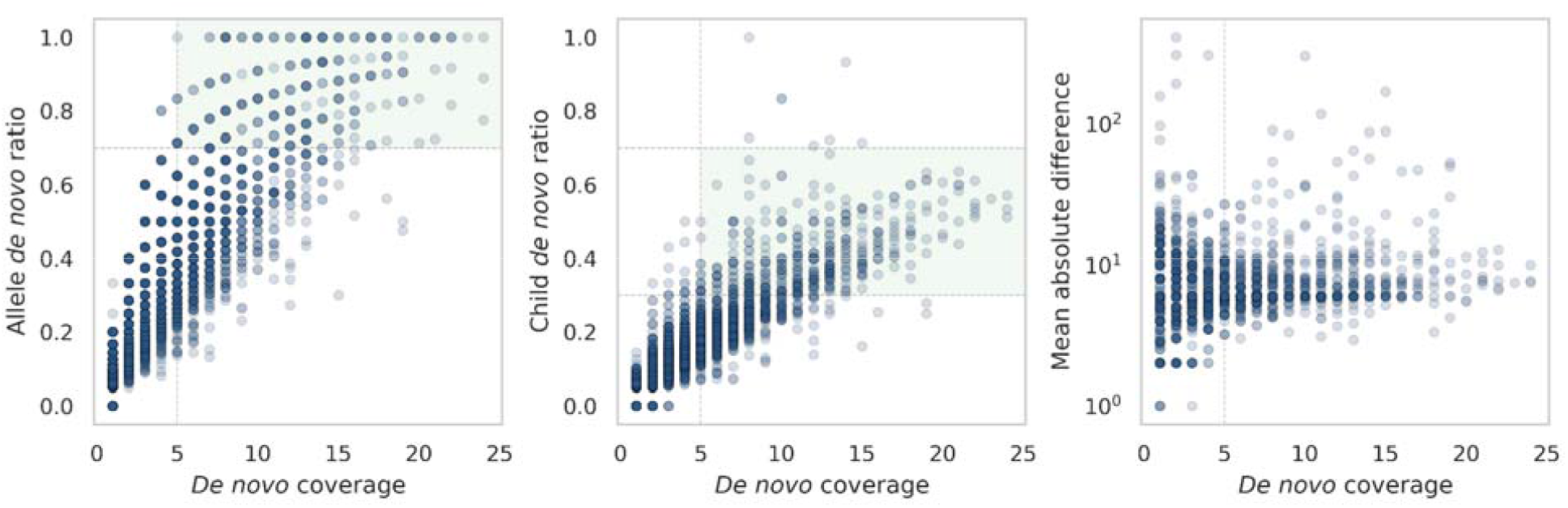
TRGT-denovo metrics. *De novo* coverage relative to the (a) allele *de novo* ratio; (b) child *de novo* ratio; (c) mean absolute difference between the reads with *de novo* evidence and P_u_. Each point represents a potential *de novo* allele. Horizontal and vertical lines indicate thresholds for minimal *de novo* coverage, allele *de novo* ratio, and a range for the child *de novo* ratio, creating shaded boxes where true *de novo* mutations are more likely.

Using these metrics, we established thresholds to identify potential *de novo* alleles (Methods). After filtering, we excluded 4,214 of the putative *de novo* sites, leaving 137 candidates. Comparison with candidates identified by the genotype-based approach using strict Mendelian consistency checks revealed an overlap of 95 candidates. This partial overlap suggests that DNMs may not always alter the allele length or may involve sub-motif changes that do not affect the motif count, such as SNVs within a repeat motif. Furthermore, small insertions, deletions, or substitutions within the repeat motifs themselves can introduce variability without altering the overall motif count. For example, a 5 bp change in a 20 bp motif is proportionally too small to affect the measured motif count. While the simple genotype-based method might miss these subtle changes, due to its reliance on counting motifs, TRGT-denovo can detect them through read-level analysis. When allowing for single repeat unit motif variations, the overlap further reduced to 29 candidates. This indicates that such small changes, typically expected in *de novo* events, were effectively excluded. In summary, TRGT-denovo outperforms genotype-based methods in detecting DNMs by capturing complex changes not reflected by simple motif counts. The simple genotype-based method identified 81,308 candidates of which only 95 (0.12%) overlapped with TRGT-denovo’s results. Indicating that TRGT-denovo effectively filters out most likely false positives. Even when the genotype-based strategy adjusts for off-by-one motif counts, it still retains 2,582 candidates out of 81,308, with only 29 (1.12%) overlapping with TRGT-denovo’s results.

### Validation of *de novo* TR calls

Previously, an in-depth analysis was performed on eight rare disease trios to identify DNMs in PacBio HiFi sequencing data, focusing on various types of variations, including TRs [10]. The study used a repeat definition catalog of 171,146 loci [11], and TRGT was used for genotyping, followed by analyzing the VCF file to detect TR DNMs. That study identified 28 *de novo* candidates, 18 of which were amenable to targeted sequencing; however, none of these putative *de novo* events were confirmed as true DNMs. Misclassifications generally occurred either because the child allele size was incorrect, or the allele was actually present in one of the parents. Using the same repeat catalog, TRGT-denovo correctly reclassified all 18 candidates as true negatives. Of the 10 unsequenced candidates, TRGT-denovo suggested that only one might be a *de novo* mutation. Note that this single unsequenced candidate was found with short read data, supporting its classification as a true *de novo* event.

To identify true *de novo* candidates in these trios, we used a larger catalog of 937,122 loci [12], and applied both TRGT and TRGT-denovo across all trios. After post-filtering (Methods), we observed 60-120 candidate DNMs per trio. A subset of these candidates was selected for validation using targeted long-range PCR and sequencing on a PacBio Sequel IIe system [10]. Based on repeat size, candidates were categorized into three groups: large (9), small (17), and those overlapping with genes associated with neurodevelopmental disorders (NDD) (6). In each category, there were 7 (large), 9 (small), and 4 (NDD overlapping) candidates for which primer design, amplification and sequencing was possible, respectively. We successfully validated 95% (19/20) of these candidate *de novo* mutations (Table 1). The one small candidate that could not be validated (shown in Supplemental Fig 1.) was a dinucleotide repeat with a single unit contraction, which, at high depth, displayed variability exceeding the *de novo* event size.

**Table 1.** Validation results. Targeted sequencing results of a subset of 20 candidate *de novo* TR calls detected by TRGT-denovo in eight trios. Calls are categorized by their size (large or small) or their genomic location, specifically in genes associated with neurodevelopmental disorders (NDD).

| Type | TR Size (bp) | TR Size $\mu$ bp (±SD) | Validated/Candidates |
| --- | --- | --- | --- |
| Large | [40, 1200] | 301 (±372) | 7/7 |
| NDD | [8, 24] | 19.25 (±6.53) | 4/4 |
| Small | [2, 5] | 3.6 (±0.94) | 8/9 |

## Discussion

The discovery of *de novo* TR mutations is highly relevant for understanding genetic disorders. However, accurately identifying these mutations presents significant challenges due to the high mutability of TRs and the vast number of repeat loci in the genome. Standard genotype filtering methods are often insufficient in this context, necessitating more sophisticated approaches. To minimize the risk of likely false positive *de novo* candidates in genotype-based *de novo* calling, an extremely high variant calling accuracy is required, reflected as a high level of Mendelian consistency (Supplemental Fig 2.). As the size of repeat catalogs grows, so does the requirement for such consistency. In this study, we introduce TRGT-denovo, a method that addresses these challenges by integrating TRGT genotyping results with read-level data from family members. We demonstrate that TRGT-denovo significantly reduces the number of *de novo* candidates compared to using raw TRGT genotypes alone. Specifically, TRGT plus TRGT-denovo identified ∼100 *de novo* TRs per trio, compared to the thousands or tens of thousands identified when using TRGT alone. This reduction in likely false positives is important for making *de novo* TR analysis more feasible and reliable in rare disease studies.

TRGT-denovo achieves high validation rates in addition to lowering the number of likely false positives. Specifically, we validated the *de novo* calls identified by TRGT-denovo, achieving a 95% validation rate (19/20 *de novo* calls). Maintaining high validation rates while reducing likely false positives not only yields a more manageable set of candidates for validation but also supports the use of larger repeat catalogs, enabling broader genomic surveys. By revisiting the genotyping-supporting sequencing data, TRGT-denovo can recapture details that might be lost in a VCF file, thereby improving the reliability of the *de novo* candidates obtained without altering the underlying genotyping process. Furthermore, we demonstrate TRGT-denovo’s utility in separate work in cases involving two trios with suspected GCC repeat expansions in the *AFF3* gene, associated with intellectual disabilities, identifying a DNM in *AFF3* as the most significant DNM across the genome, further substantiating its pathogenic significance [13].

Although TRGT-denovo significantly reduces (likely) false positives in trio analysis, there are still areas for improvement. *De novo* assessment of TRs remains notably challenging, despite the advancements enabled by long-read sequencing and TRGT-denovo. Current work aims to introduce haplotype matching across samples to enhance the reliability of inheritance inference, further reducing false positives by addressing issues like parental allele dropout. TRGT-denovo enables more accurate TR mutation studies, potentially leading to new insights in genetic disorders and genome dynamics.

## Methods

### TRGT-denovo

TRGT-denovo is a method for identifying DNMs within TR loci. It is designed to work in tandem with TRGT, a tool for targeted TR genotyping using PacBio HiFi sequencing data. TRGT requires aligned HiFi reads and a set of repeat definitions (Fig. 1a, b). The generated outputs include a VCF file—with full-length repeat consensus allele sequences and genotyping information—and a BAMlet file with segments of HiFi reads spanning each repeat allele. TRGT also integrates haplotype phasing tags in the BAM from tools such as HiPhase [14] or WhatsHap [15], in addition to using mismatches surrounding the TRs for allele phasing. TRGT-denovo analyzes TRGT VCF and BAMlets from family trios. Each repeat locus is considered independently, requiring successful genotyping from each family member, proceeding as follows:

### Within-sample read partitioning

TRGT-denovo extracts reads spanning each repeat allele in each family member for a specific locus of interest (Fig. 1c.). For each allele a consensus sequence is generated, consisting of the TRGT derived consensus allele sequence and 50bp of flanking genome sequence. Typically, with two alleles per family member, this results in six of such sequences per trio, representing the alleles of the child (C_0_, C_1_) and those of the parents (F_0_, F_1_ from the father, and M_0_, M_1_ from the mother). Subsequently, within each family member, the reads are partitioned to their corresponding alleles using the TRGT-obtained allele assignments by default (Fig. 1d.). Partitioning may optionally be based on the read’s alignment to consensus sequences or available phasing data. When partitioning relies on alignment, reads are aligned to their sample-specific consensus sequences through end-to-end, gap-affine, two-piece alignment, with assignment to the consensus sequence with the highest alignment score. Ties in scores are resolved by random selection, phasing data, or TRGT allele assignment. If phasing is used for partitioning, reads are assigned according to their allele based on phasing tags; in the absence of such tags, alignment as previously described serves as the fallback method. Alignment plays a fundamental role in capturing the inherent variation across the reads. This is true regardless of the partitioning strategy used; for each allele, the alignment scores of each read are always obtained from all reads assigned to that allele. These scores indicate how closely the reads resemble the allele and reflect the distribution of uncertainty across all assigned reads within the alleles of each family member. To efficiently manage millions of alignments of highly similar sequences, we have used the Wavefront alignment algorithm (WFA) [16], which exploits similarities between sequences to accelerate the computation of the optimal alignment.

### Candidate *de novo* allele alignment and analysis

Each child allele is evaluated as a potential *de novo* allele by examining its similarity to the parental alleles. This is achieved by aligning each read (previously partitioned into the alleles F_0_, F_1_, M_0_, M_1_, C_0_, and C_1_) against the child’s allele consensus sequences, generating alignment score distributions for reads relative to the child alleles C_0_ and C_1_, as shown in Fig. 4.

**Fig 4.**
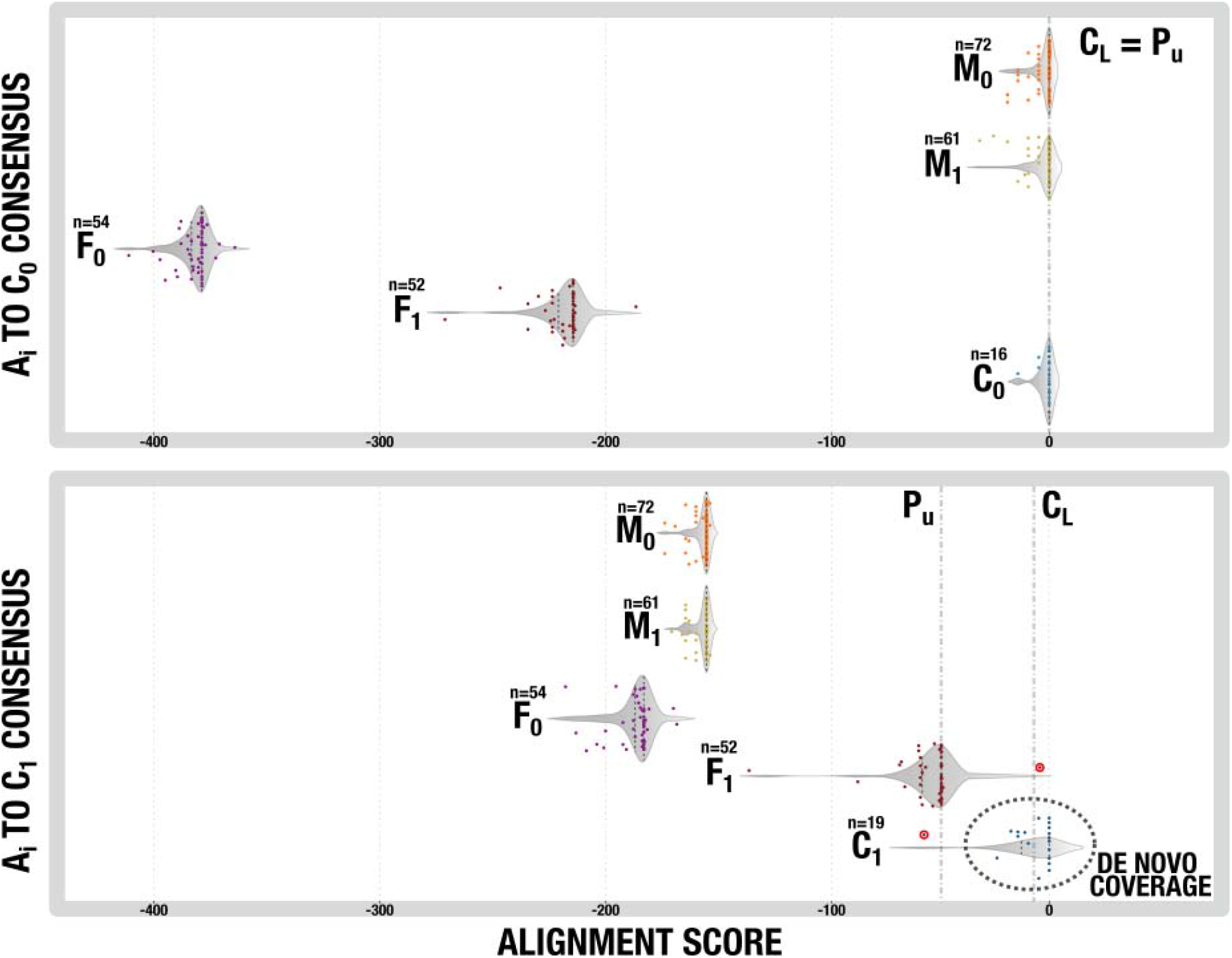
Alignment score distributions. Distributions of alignment scores for reads spanning alleles (M_0_, M_1_, F_0_, F_1_, C_0_, C_1_) when aligned to alleles C_0_ (a) and C_1_ (b). WFA alignment scores range from negative, less similar, to zero, perfect match. Inheritance patterns, as inferred from surrounding genetic variation, are: M_0_ → C_0_ (inherited) and F_1_ → C_1_ (inherited + *de novo*). Symbols P_U_ and C_L_ denote the parental upper bound and candidate *de novo* allele lower bound respectively. Each point corresponds to an individual read aligned to C_0_ or C_1_. Red-outlined points highlight two observations: a read from F_1_, exceeding C_L_, showing overlap with C_1_, and a read from C_1_ that falls below P_U_, and overlaps with F_1_. In C_1_ there are 19 reads, of which 18 exceed P_U_, contributing to the *de novo coverage*.

Alignment score distributions, denoted by 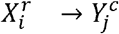 for some *i, j*, reflect the scores from aligning reads of allele *X*_*i*_, 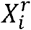, to the consensus sequence of allele *Y*_*i*_, 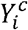. For instance, 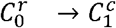 represents the scores of reads in allele C_0_ when aligned to allele C_1_. Self-0 1 alignments, such as 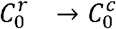 and 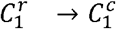 typically show maximum WFA alignment scores (zero scores), with deviations recorded as negative values. An exception being mosaic alleles, like some FMR1 expansions, where numerous reads differ significantly from the consensus sequence. In instances where a child inherits both alleles, the majority of the 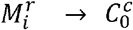 distribution is expected to overlap with 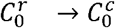 and 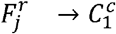 with 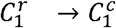 for some *i, j* or vice versa (Fig. 4a.). Conversely, if allele C_1_ is *de novo*, the 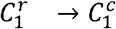 distribution will separate, diverging from either 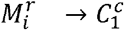 or 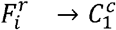 for all *i* (Fig. 4b.). Note that this divergence is always unidirectional; for instance, if C_1_ is *de novo* relative to F_1_, then 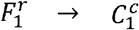 will shift left relative to 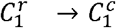. Absence of such a shift would imply no underlying difference and, consequently, no DNM. Note that any divergence, regardless of changes in sequence length or composition, can signal a DNM and will cause a corresponding shift in the distribution.

Identification of DNMs relies on analyzing shifts in alignment score distributions, based on the assumption that the closest matching parental allele distribution to the child’s allele usually reflects an inherited allele without mutation. Significant shifts in these distributions suggest the presence of a DNM, indicating that the allele, although likely derived from the closest parental allele, has undergone changes and is not inherited as is. A significant divergence in distributions suggests greater dissimilarity, which acts as a measure of the magnitude of DNMs, enabling differentiation based on the size of these mutations. These distributions also facilitate estimates of inheritance patterns, where reads from an inherited allele are expected to closely match those of the corresponding child allele, though this can be complicated by scenarios such as multiple identical parental alleles. To quantify divergence, a parental upper bound, P_U_, is defined, derived from the 1.0 quantile of the parental distribution, and is used as a reference point against which all child allele reads are tested. Reads with scores exceeding P_U_ indicate a potential DNM, shown as *de novo coverage* in Fig. 4b., with nearly all C_1_ reads surpassing this threshold. Conversely, comparing parental read scores relative to the child allele’s lower median bound, C_L_, reveals shared traits. This *overlap coverage* aids in differentiating characteristics between parents and child. For example, overlap in reads, such as seen in 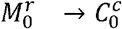 and 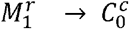 with 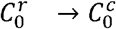, indicates inheritance of C_0_ without mutation. In contrast, C_1_ represents a DNM, as demonstrated by the lack of overlap in nearly all parental allele distributions 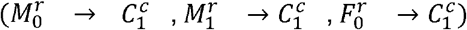 with 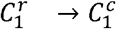. An exception being the single read showing overlap in 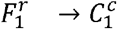; however, this is proportionally negligible against the background of all other reads in 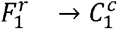 that do not overlap with 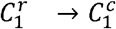. The magnitude of DNMs is further quantified by calculating the mean absolute difference between the *de novo* reads and the P_U_ threshold. In cases of inheritance (Fig. 4a.), this value is zero, reflecting (near)-perfect alignment between parental and child alleles, with any deviation denoting the presence and scale of DNMs.

### Detecting *de novo* TR mutations

TRGT-denovo measures and outputs metrics for every single child allele, generating metrics such as the *de novo* coverage, overlap coverage, and the magnitude of potential DNMs. Note that the *de novo* and overlap coverage should be considered in the context with the allele-specific and total coverage at the locus for each family member i.e., their ratios. For example, as shown in Fig. 4b. C_1_ has 19 reads, with 18 exceeding the P_U_ threshold, yielding an allele *de novo* ratio of ∼0.95. A high *de novo* ratio strongly suggests a DNM, assuming sufficient overall coverage. Conversely, a low ratio may indicate weaker evidence or suggest complexities like segmental duplications, mosaicism, or stutter. *De novo* coverage is a strong predictor for detecting *de novo* variants, correlating with increased allele *de novo* ratios, approaching 1.0, and child *de novo* ratios converging to 0.5 (Fig. 3). This indicates that higher coverage helps in accurately identifying true *de novo* mutations by ensuring that these ratios align more closely with the expected balance. Note that all loci should be compared against the entire set of tested loci, creating a distribution that aids in defining thresholds for identifying likely DNMs. This comparative approach also helps to identify coverage imbalances or sample mix-ups, ensuring reliable results. Based on empirical data, the following filtering criteria have been established as a starting point:

1. Minimum *de novo* coverage of 5: This threshold ensures enough read support to confidently call a DNM. Lower coverage may result in insufficient evidence to distinguish true mutations from sequencing noise or errors.
2. Allele *de novo* ratio of at least 0.7: A high allele *de novo* ratio indicates that the majority of reads for an allele exceed the parental upper bound threshold. Observations show that true DNMs exhibit a strong divergence from parental alleles, and ratios below 0.7 often suggest weaker evidence or potential artifacts.
3. Child *de novo* ratio between 0.3 and 0.7: This range is chosen to balance the need to detect *de novo* events while accounting for the possibility of allelic dropout or partial mosaicism. Ratios below 0.3 or above 0.7 could indicate sample anomalies or errors in read assignment.
4. Low probability for allelic dropout: we require at least one read supporting each haplotype, as obtained from available within-sample phasing. This helps to avoid likely false positives due to incomplete haplotype representation.

These criteria were determined through a combination of empirical testing on known *de novo* mutations and analysis of genomic datasets. However, they should be considered as guidelines rather than strict rules and may need to be adjusted based on specific study designs or sequencing depths.

## Competing interests

T. Mokveld, E. Dolzhenko, Z. Kronenberg, and M. A. Eberle are employees and shareholders of Pacific Biosciences.

## Funding

T. J. Nicholas, T. Sasani, and A. Quinlan were supported by the following awards from the National Institutes of Health: RC2TR004391 and R01HG010757. H. Dashnow was supported by the following award from the National Institutes of Health: K99HG012796

## Notes

### Competing Interest Statement

T.M., E.D., Z.K, and M.A.E are employees and shareholders of Pacific Biosciences.

