## Supplemental figures for "TRGT-denovo: accurate detection of *de novo* tandem repeat mutations"

### Supplemental materials

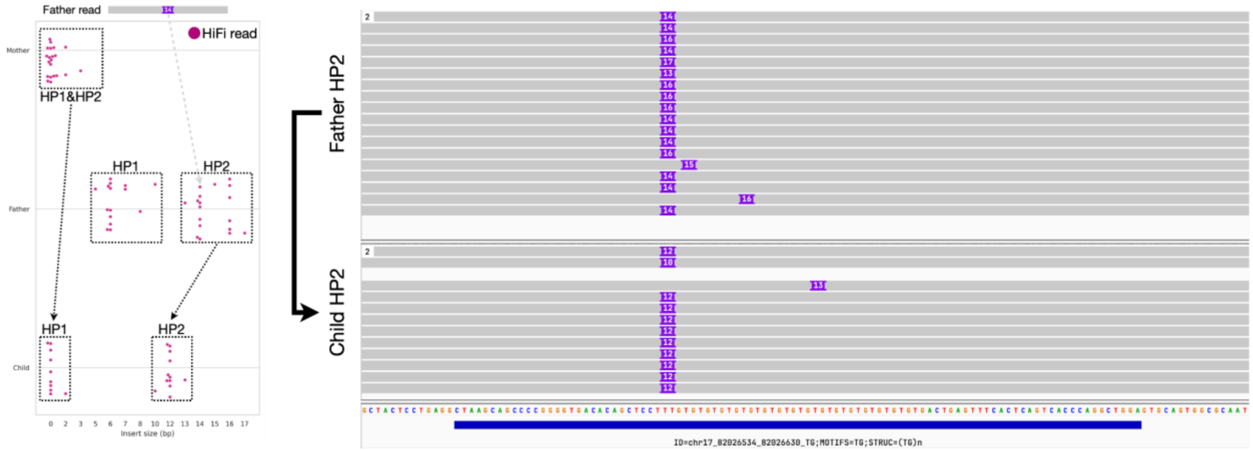


**Supplemental Fig 1. False positive *de novo* call**. The candidate *de novo* call that could not be validated with targeted sequencing, a single unit contraction inherited in child haplotype 2 from father haplotype 2 (as derived from surrounding variation). Left: for each family member the insertion size is extracted from the read data inside the TR locus and plotted. Right: IGV plot of the relevant haplotypes in father and child.

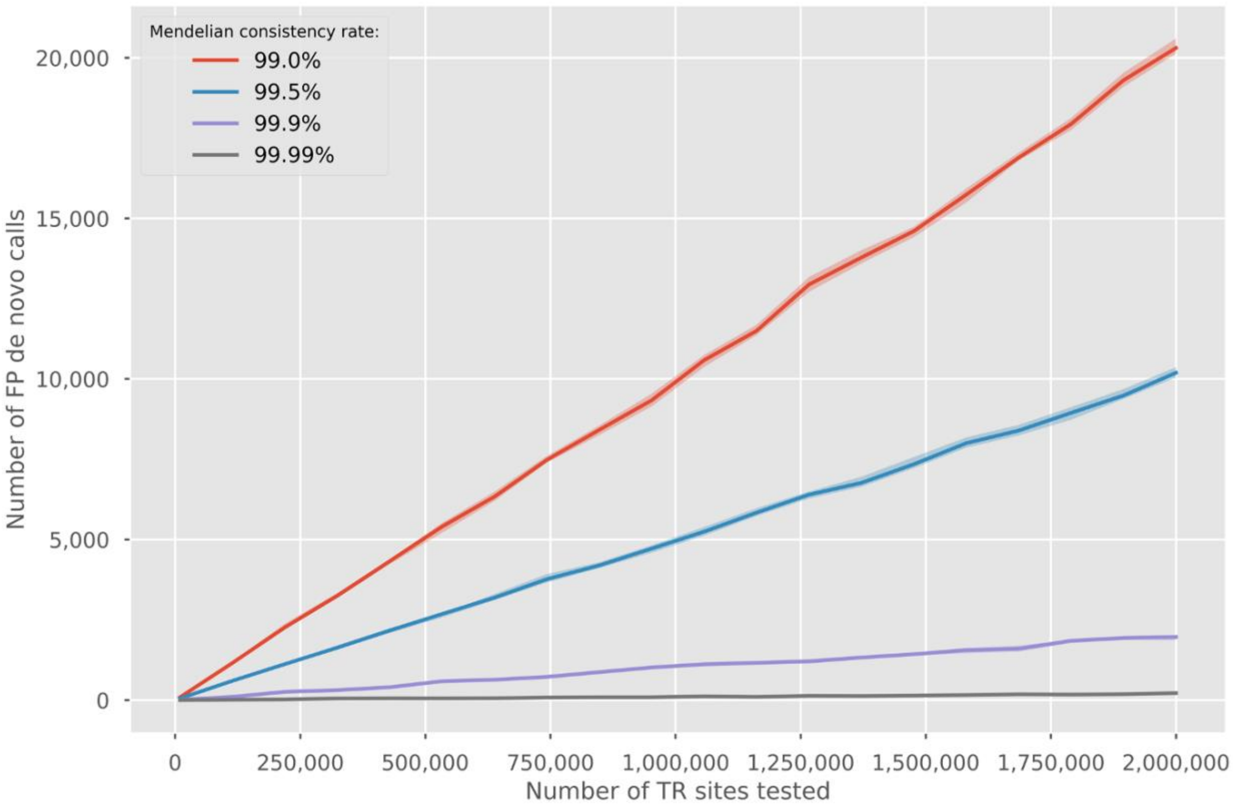


**Supplemental Fig 2.** The inevitability of false positive *de novo* TR calls when the number of tested TR loci increases, given different rates of mendelian consistency.
